# Genetic evidence suggests relationship between contemporary Bulgarian population and Iron Age steppe dwellers from Pontic-Caspian steppe

**DOI:** 10.1101/687384

**Authors:** Svetoslav Stamov, Todor Chobanov

## Abstract

Ancient DNA analysis on the ancestry of European populations conducted in the last decade came to the puzzling conclusion that while all contemporary European populations can be best represented as an admixture of 3 ancestral populations –Early European Neolithic farmers (ENF), Western Hunter-Gatherers (WHG) and Ancestral North Eurasians (ANE), contemporary Bulgarians and few other SEE populations can also be represented as an admixture of two groups only – Early European Neolithic farmers and contemporary Caucasian people equally well.

If modeled as an admixture of two groups only, the ANE component presented in contemporary Bulgarians would have arrived on the Balkans with hypothetical ANE (Ancestral North Eurasians)-rich Caucasian population.

In this paper, we test the hypothesis that increased Caucasian component in contemporary SE Europeans, has been introduced on the Balkans by migrating Iron Age steppe dwellers from Pontic-Caspian steppe. We analyze available DNA datasets from both ancient and contemporary samples and identify a Caucasian signal, carried to Balkan populations by the nomadic dwellers of Early Medieval Saltovo-Mayaki Culture, located on the northern slope of Caucasus Mountains and adjacent steppe regions. We also identify two additional sources of Caucasian admixture in SEE populations, which are not specific to Bulgarian population only. Based on the results from our population genetic analysis we suggest that contemporary Bulgarians are an admixture of ancestral Slavonic groups, rich on locally absorbed EEF DNA and Proto Bulgarians, rich on Caucasian DNA and genetically related to the bearers of the Saltovo-Mayaki Culture from 8-10 century AD.

## Introduction

All contemporary European populations can be represented as an admixture of 3 ancient groups: Early European Neolithic farmers (ENF), western hunter-gatherers (WHG) and Ancestral North Eurasians (ANE). (Lazaridis I, Patterson N, Mittnik A, et al. Ancient human genomes suggest three ancestral populations for present-day Europeans. *Nature*. 2014;513(7518):409-13.)

**Fig. 1.**
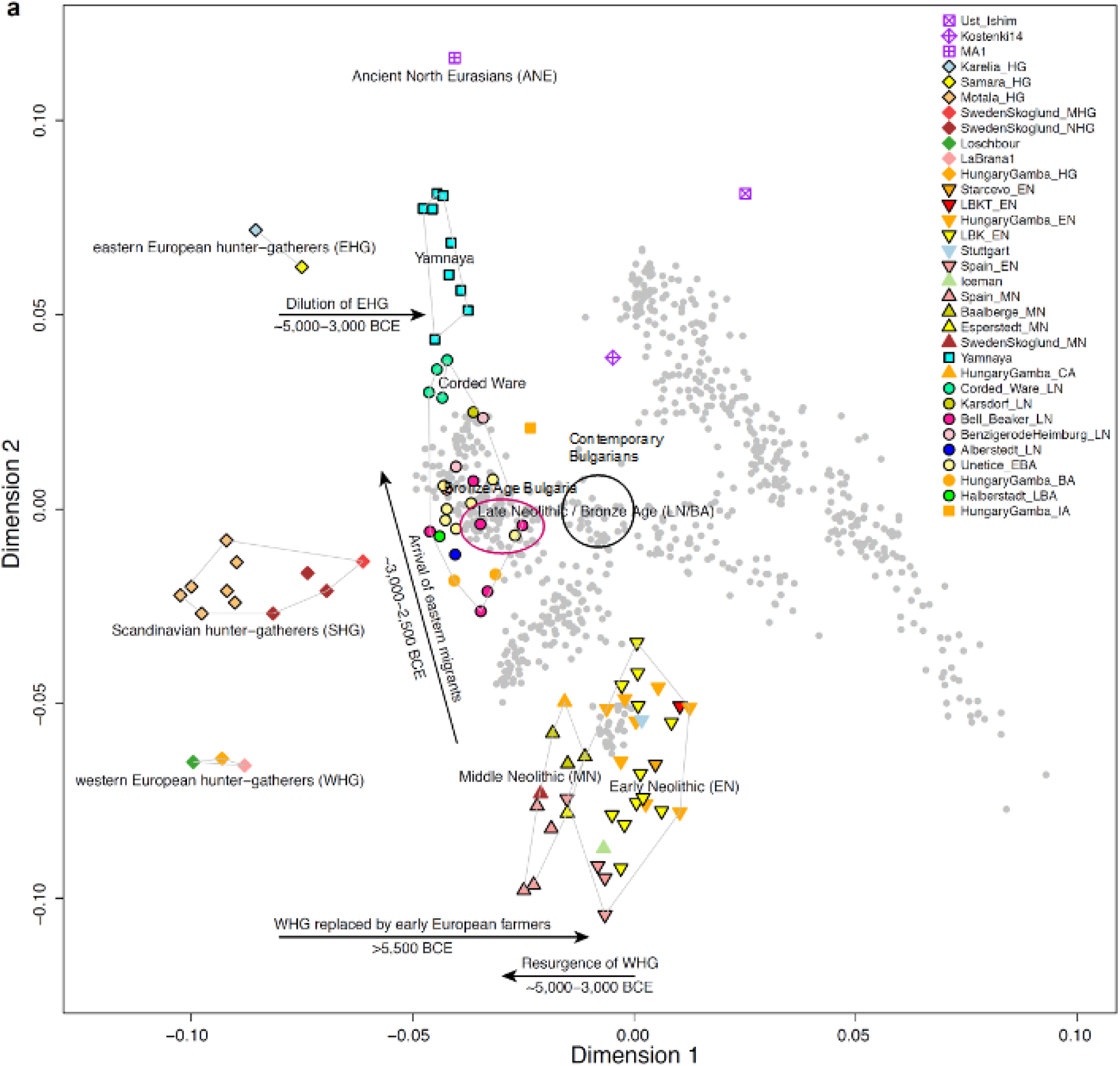
Contemporary Bulgarians show an extra layer of Caucasian admixture, which is missing from the Bronze Age Balkan population (BAB). BAB are a mixture of Yamna migrants and EEF - just as rest of European populations. On the plot we can see that contemporary Bulgarians are closer to the Caucasian cluster than Bronze Age Balkan samples are. PCA after Haak W, Lazaridis I, Patterson N, et al. Massive migration from the steppe was a source for Indo-European languages in Europe. Nature. 2015;522(7555):207-11., Mathieson I, Alpaslan-Roodenberg S, Posth C, et al. The genomic history of southeastern Europe. Nature. 2018;555(7695):197-203

On the map (Fig. 1) contemporary Bulgarians are distributed nearer to contemporary Caucasians than most European populations which suggests an extra degree of Caucasian admixture that has been absent in the rest of Europe. This implies admixture events that are specific to Bulgarian population and whose effects are limited to the area of Balkan Peninsula mostly.

Laazridis and Reich first noted that like the rest of Europeans, south-east Europeans can best be modeled as 3-way admixture (ANE-EEF-WHG), however they can be modeled as 2-way only admixture equally well (EEF-Caucasians, where ANE component would have come from additional Caucasian migrations to the Balkans). Haak et al confirmed the findings of D. Reich and established a vector of massive migration from Black Sea – Caspian steppe region into Europe. This migration occurred during early Bronze Age and became major contributing factor to the populations of all contemporary Europeans. Haak established that BA migrants represented an admixture of Caucasian Hunter Gatherers, genetically rooted in Mesolithic Northern Iran and East European Hunter Gatherers from what is now Russian plain. The migrants carried distinctive Caucasian signature and introduced Caucasian component throughout European continent. While this signature had been dilated in Western Europe in the centuries that followed, it had increased in the Balkan populations. This increase is suggestive of more admixture events with populations, caring Caucasian component and limited to the Balkans only. (see Fig. 1.)

The increase in Caucasian component in contemporary Bulgarians postdates Bronze Age migrations. Historical literature suggests that the arrival of this component in Bulgarian population could be related to the migration of Protobulgarians (Bulgars) during 6-8 century AD and the foundation of First Bulgarian Kingdom (V. Zlatarski, S. Runsiman, R. Rashev). Century long archeological research has identified northern Caucasian slopes and adjacent Kuban River zone as the likely homeland of the migrating Bulgars. Archaeological research suggests intensive contacts between Bulgars and the neighboring Caucasian and Alanic tribes, including the emergency of material culture of mixed origin, suggestive of a synthesis between IA Caucasian and IA steppe traditions, emerging in the zone of Cuban river during Saltovo-Mayaki Culture (SMC, 8-10 century AD). In this paper, we present the results from our analysis on the available ancient genetic data from BA and IA Western Eurasia, including samples from SMC in their relation to modern Bulgarians.

## Method

We analyzed ancient DNA samples from Bronze Age, Iron Age and medieval Western and Central Eurasia. In an attempt to establish the source population and the timing of the additional Caucasian admixture in contemporary Bulgarians, we merged the ancient dataset with the dataset of 40 contemporary Bulgarians as well as the dataset of 100 contemporary individuals from neighboring populations. We computed principal component analysis on the present populations and projected available ancient DNA samples from Western and Central Eurasia. We also built a neighbor joining tree of the available ancient and contemporary samples. All genetic trees and PCA plots have been computed with PAST software for palaeogenetic DNA analysis.

We also reviewed already published genetic research on the topic in the scientific literature in order to identify what has been already known about the timing and the hypothesized source population. We also test several well-known historical hypothesizes about the origins of contemporary Bulgarians and early IA Protobulgarians.

## Results

Using statistical genome-wide analysis, we detected nontrivial genetic connection between contemporary Bulgarians, inhabitants of Bronze Age Armenian plateau and Iron Age dwellers from SMC. Our analysis also suggests surprising connection between contemporary Bulgarians and Iron Age Scythians from Hungarian plain.

### Principal Component Analysis

For our PCA and genome-wide statistical analysis we used PAST3.22, version December 2018 - Paleontological statistics software package for education and data analysis (Hammer 2001).

All contemporary individual DNA genome-wide data files were retrieved from Yunusbayev et al 2012. To analyze the genetic distances and genetic relationship of the retrieved samples to the contemporary Bulgarian samples, we built several principle component analysis (PCA) plots, which visualized the genetic relationship between the individuals, their genetic contribution to the contemporary Bulgarians and we created several genetic trees based on their degree of relatedness.

In our first PCA (Fig 2) we combined dataset from 137 ancient samples from the Eurasian Steppe - from what is now Mongolia to what is now Hungarian plain (P. Damgaard et al, Nature volume 557, pp369–374, May 2018) and merged it with selected contemporary individuals from SE Europe (dataset from Yunusbayev et al 2012)

**Fig. 2.**
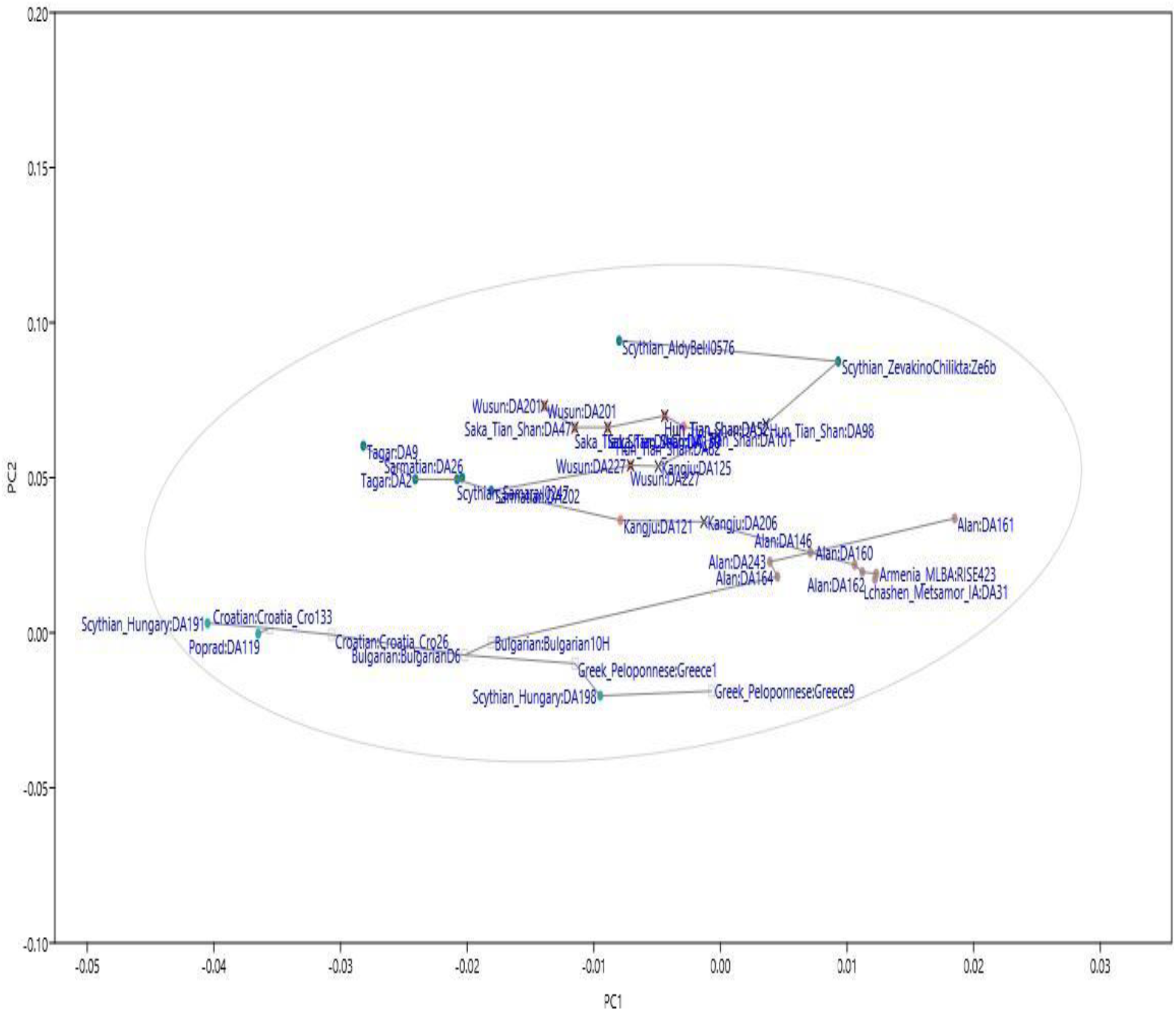
PCA on the relationship between contemporary Bulgarians and ancient samples from BA and IA Eurasian steppe. While none of the contemporary Bulgarians yields relation to the ancient CA populations, PCA1 suggests genetic connection between contemporary Bulgarians and IA individuals *AlanDA243, AlanDA164 and Alan DA146* from North Ossetia and SMC.

The results of PCA (Fig 2) renders direct connection between contemporary Bulgarians and Inner Asian steppe nomads from migration period unlikely. None of the contemporary Bulgarians yielded any direct or mediated relation to the ancient Far Eastern and Central Asian nomadic steppe populations.

In order to examine population transformation in what is now contemporary Bulgaria from early Bronze Age trough Iron Age till now, we also added 8 ancient samples from the late Neolithic / Early Bronze Age and early Iron Age, which we retrieved from Haak et al 2015, 207-11 and from Mathieson et al 2018, 197-203.). We present the results in Fig. 3

**Fig 3.**
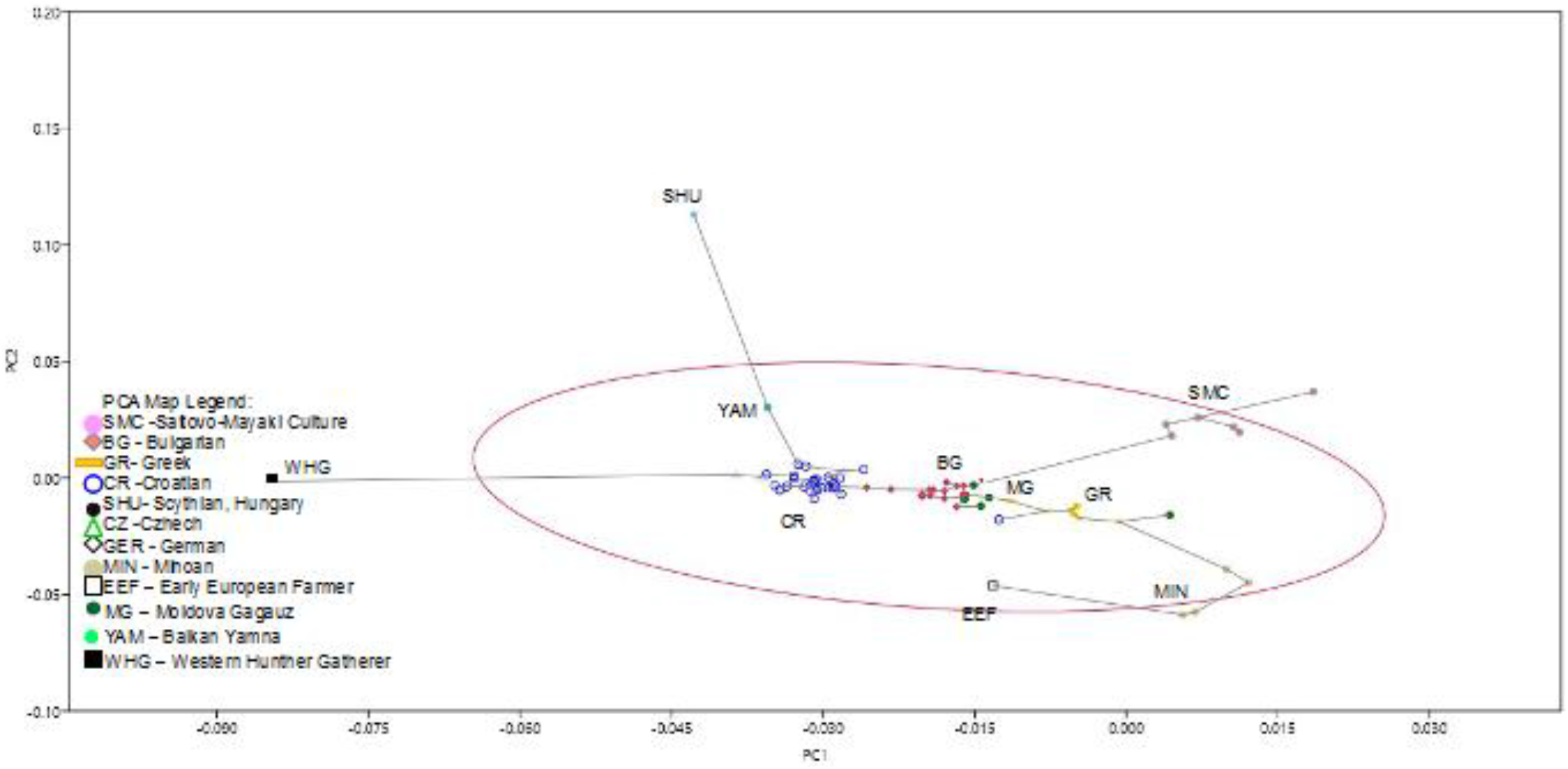
There is statistically significant relationship between contemporary Bulgarians and the Protobulgarians from SM. The genetic affinities detected by PAST3 suggest that SM people have contributed to contemporary Bulgarians only and their contribution to the rest of Balkan population has been transmitted from contemporary Bulgarians to their geographical neighbors.

PCA results suggest genetic connection between contemporary Bulgarians and the ancient individuals AlanDA243, AlanDA164 and Alan DA146 belonging to SM culture.

In our next PCA we added Scythian samples from Hungarian plain from 4^th^ Century BC (classical antiquity). The plot suggests connection between Scythian samples, European Alans from the migration period and the nomads from the *Saltovo-Mayaki* Culture as all 3 groups showed genetic connection to contemporary Bulgarians. (fig. 4)

**Fig. 4.**
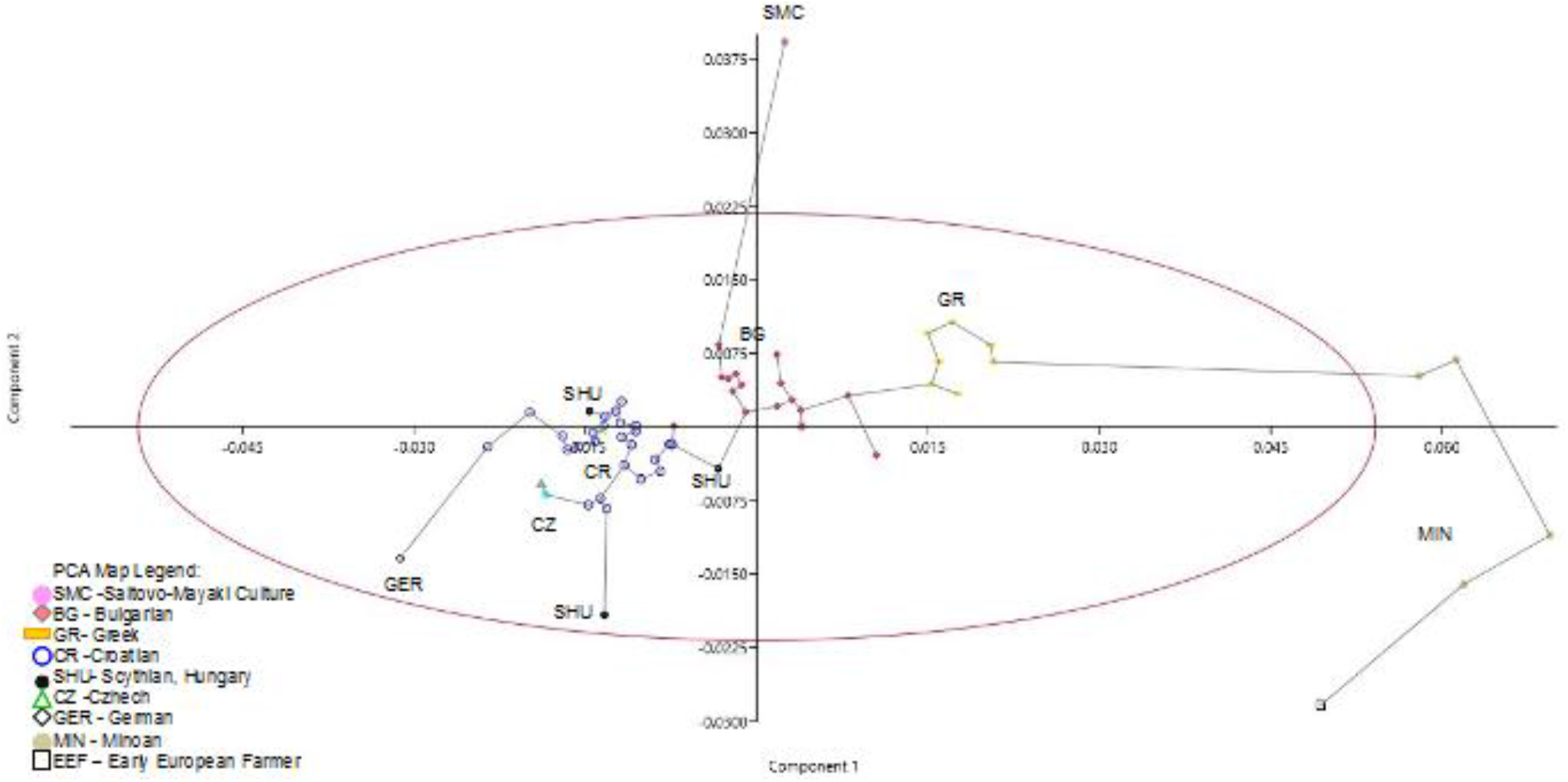
These results imply nomadic influence from migration period being carried over to the population genomics of contemporary Bulgarians.

Our PCA (Fig.2) also revealed indirect connection between contemporary Bulgarians and central Asian Bronze Age nomads of East Iranic origin known as Kangju group. This relation however is dependent on the presence of sample Alan DA146 from *Saltovo-Mayaki (Saltovo*, SM for short) culture on the PCA Plot and disappears if we remove this sample from the plot. We suggest that this discrete connection represents earlier stages of the migration of certain proto SM groups (Sarmatians-Alans?). Yet the rest of SM samples did not yield same connection to Kangju but showed detectable connection to the samples from Bronze Age Armenian plateau (fig. 2), suggestive of multiple admixture events during different earlier stages of migrations and contacts of SM people, as one of these stages must have included Armenian plateau in Central Caucasus.

Since there were multiple waves of migration from Caucasus to the Balkans including IE migration during Bronze Age and the emergence of Minoans during early BA and they all carried substantial Caucasian component with them (Haak et al 2015,207-11. and Mathieson et al 2018, 197-203), in our next plot we tried to distinguish the admixture signal coming from SM people from admixture signals coming from the earlier migrations. In order the test the Huns as potential carriers of the same signal, we also included a sample of iron-age Siberian hunter gatherer as a proxy for the Huns and in order to test the early Slavs for yet another potential carrier, we included contemporary Croatian samples as a proxy for the medieval Bulgarian Slavs. We also included Moldova Gagauz samples to test if they carry stronger Protobulgarian signal as it has been hypothesized by some of Bulgarian historians. We present the results in Fig. 5 and Fig. 6:

**Fig. 5.**
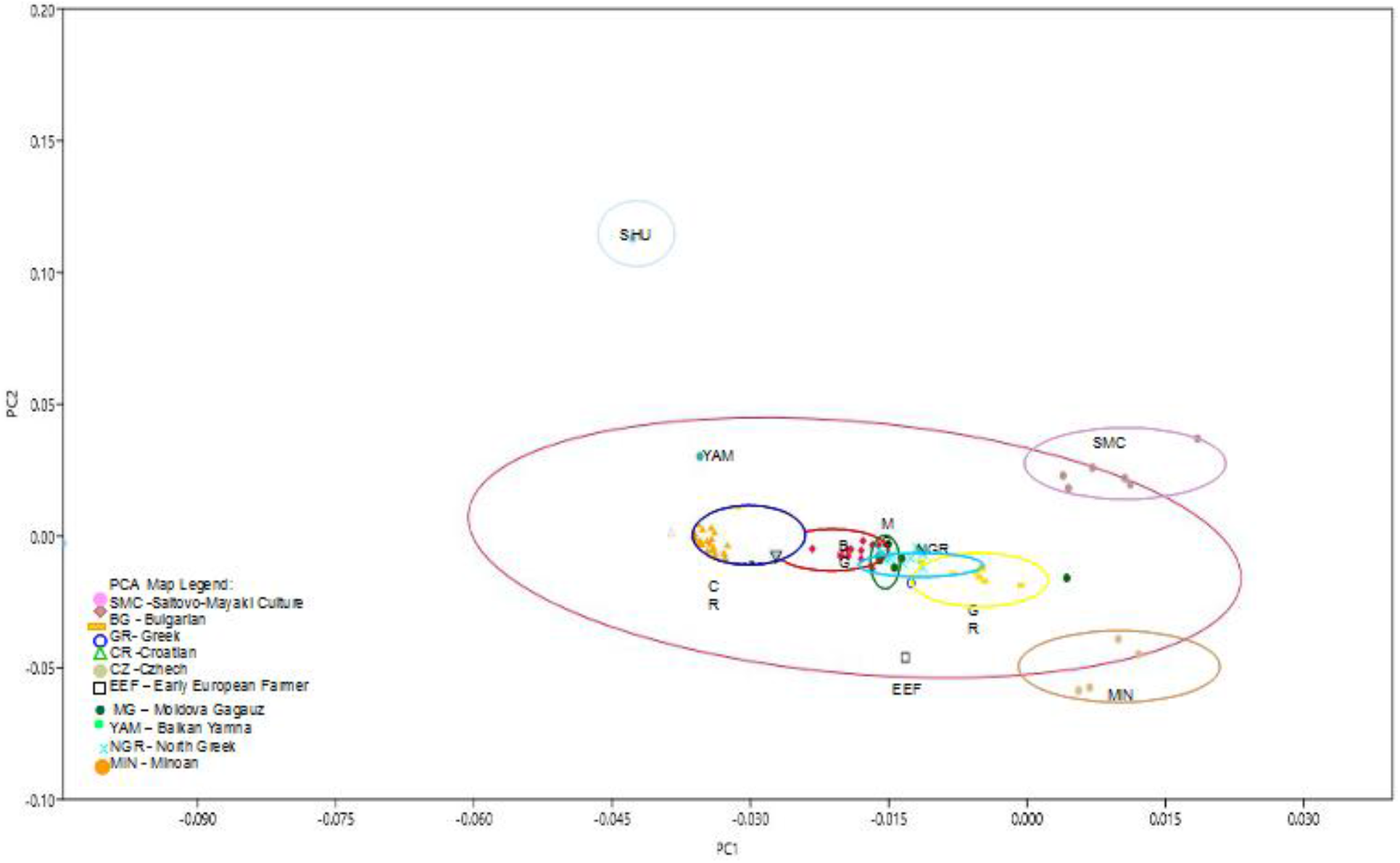
In the PCA plot (fig. 5) the current Balkan nations form a cline. None of the tested samples showed detectable relation to SHG sample. The signals coming from SMC, NPBA Minoans and Bronze Age proto Thracians are clearly distinguishable from each other. Moldova Gagauz samples take intermediary position between contemporary Bulgarians and Contemporary Greeks and do not show stronger connection to SMC than contemporary Bulgarians, hence the signal from Protobulgarians in contemporary Bulgarians comes directly from SM and is not mediated by Gagauz people (which also carry this signal). Bronze Age proto Thracians are genetically closer to early medieval Slavs (represented here by Croatian samples) than to contemporary Bulgarians and their influence on Bulgarian population genomics is not direct, but is probably mediated by early Slavs;

Peloponnese Greeks show closest affinity to Neolithic Peloponnesus and Bronze Age Minoans (fig. 5 and fig. 6). We conclude that the influence of Minoans on contemporary Bulgarian population is not direct and is due to population transfers and exchanges that led to admixture between medieval Bulgarians, medieval Greeks and medieval ERE populations. Both contemporary Greeks and contemporary Bulgarians show considerable distance to Bronze Age Balkan Yamna population (Thracians?) and Thracian contribution is mediated by the Croatians (fig. 5) as a proxy of the early Slavs, unless it masks Illyrian contribution in contemporary Croatians. We cannot determine whether Croatian samples reflect Illyrian or Thracian influence on the genomes of early Slavs based on the available data only. Further research is needed to clarify this topic.

We noted that SM (Protobulgarian-Alan) influence among contemporary Balkan nations has its strongest representation in contemporary Bulgarians (Fig 4) where it arrives directly and this Protobulgarian influence in the other Balkan nations is mediated by the contemporary Bulgarians who channel it.

Neighbor joining tree, built with PASTX software on the base of genetic relationship between the samples:

**FIG. 6.**
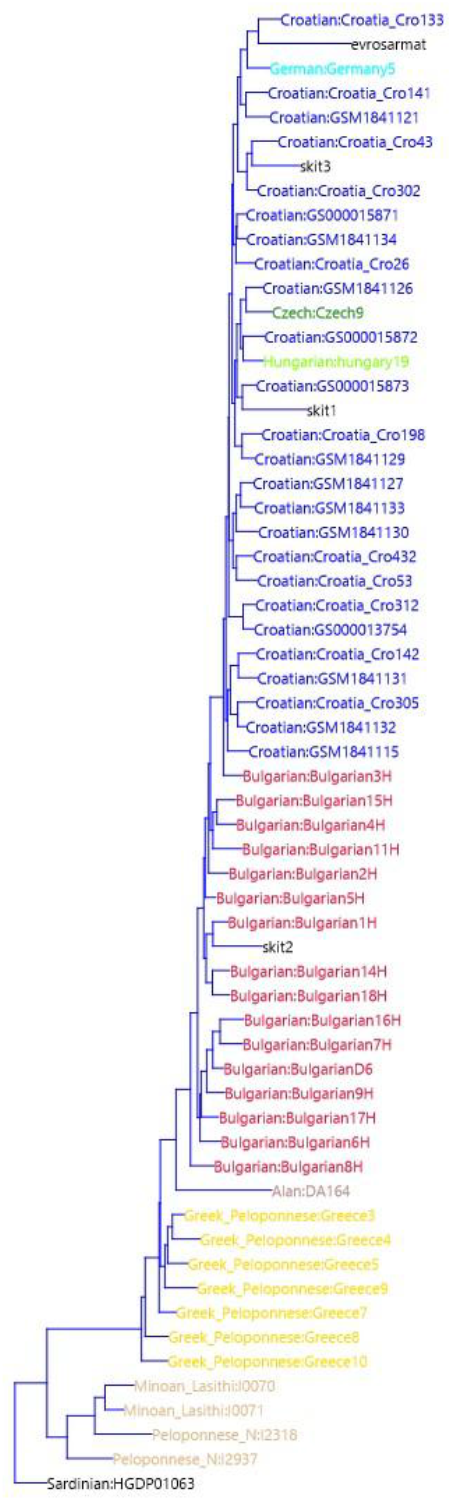
Neighbor joining tree

## Conclusions from the DNA data analysis

The results suggest that SMC related populations are among the precursor of contemporary Bulgarians. This makes SM culture at its precursor stage (600-700 AD) leading candidate for the source population of Asparukh Bulgarians. These results also suggest that Asparukh’s tribe(s) are indistinguishable from the Sarmato-Alanic groups from Early MA and Late antiquity and, surprisingly, do not carry Siberian and Central Asian admixture on the Balkans with them. Unlike BA Thracians and the early Slavs, SMC carry substantial Caucasus admixture, related to the tribes from Bronze Age Armenian plateau and seems to have transmitted this admixture to the contemporary Bulgarians. The relationship between Protobulgarians and Sarmato-Alanic tribes from the Late antiquity and Early medieval epoch remains to be clarified further, however genome wide-data suggest that Protobulgarians were themselves an admixture in equal proportions between two close, but distinct populations −1. Alano-Sarmatian tribe from the region north of Caucasus with some Kangju link to it and 2. Unknown tribe(s) originating from what is now Armenian Plateau. Both Scythian samples from the Hungarian steppe and the Alans from *Saltovo-Mayaki* culture bear strong genetic resemblance to the Bronze Age Caucasian samples, which is missing in central Asian nomads but is presented in the contemporary Bulgarians.

Our results cast a doubt on a connection between Inner Asian nomadic tribes from Antiquity and the Protobulgarians-Alans from SM culture and Northern Caucasus. The lack of Inner Asia autosomal DNA links for the Protobulgarians confirms the results from the mtDNA sampling of materials from 8^th^-9^th^ c. necropolises on the Lower Danube. The main haplogroup H (H, H1, H5, and H13) prevalent in European populations has a 41.9% frequency in modern Bulgarians, and it was observed in 7 of 13 proto-Bulgarian samples. Again no evidence was found of East Asian (F, B, P, A, S, O, Y, or M derivative) haplogroups (Nesheva et al 2015, 22). An earlier major representative survey of present dale male lineages in Bulgaria (over 800 individuals) revealed that *“Haplogroups C, N and Q, distinctive for Altaic and Central Asian populations, occur at the negligible frequency of only 1.5%.”* (Karachanak et al 2013). Our research suggest that author’s conclusion of the survey that *“…our data suggest that a common paternal ancestry between the proto-Bulgarians and the Altaic and Central Asian populations either did not exist or was negligible…”*(Karachanak et al 2013, abstract) was correct.

Since the debate about potentially “autochthonous” component in the contemporary Bulgarians (present day version of “Illyrism”) has become somewhat hotly debated topic in Bulgarian society today, we also clarified the origin of this Caucasian component further and managed to split the Caucasian component coming from SM from the Caucasian components already presented on the Balkans prior to Protobulgarian migration. We established that while all three carry somewhat similar Caucasian component (fig.3, fig.4, fig.5), the signal, coming from SM is the strongest in contemporary Bulgarians, the signal coming from Bronze Age Thracians is the strongest in contemporary Croatians and the signal, coming from Bronze Age Minoans is the strongest in contemporary Greeks. These three signals clearly differ from each other and their source populations are clearly distinguishable. Yet all tree carry an excessive Caucasian component, suggesting non-local origins for all three of them and suggestive of at least three different migrations from the Caucasus Mountains and adjacent regions to the Balkans. However, contemporary Bulgarians have received their Minoan component mostly through population exchange with Byzantium and their Bronze age Thracian component trough admixture/population exchange with early medieval Slavs and Croats. The signal that distinguished contemporary Bulgarians from the other Balkan nations is the unique signature of SM-Alan people, who appear amongst the direct precursors of contemporary Bulgarians.

## Supplement

Archaeological overview on the formation of Asparukh’s Protobulgarians.

## References

1. Lazaridis I, Patterson N, Mittnik A, et al. Ancient human genomes suggest three ancestral populations for present-day Europeans. Nature. 2014;513(7518):409–13.

2. Haak W, Lazaridis I, Patterson N, et al. Massive migration from the steppe was a source for Indo-European languages in Europe. Nature. 2015;522(7555):207–11

3. Mathieson I, Alpaslan-Roodenberg S, Posth C, et al. The genomic history of southeastern Europe. Nature. 2018;555(7695):197–203

4. P. Damgaard et al, 137 ancient human genomes from across the Eurasian steppes, Nature, Nature Springer, May 9 2018

5. Yunusbayev et al. The Caucasus as an asymmetric semipermeable barrier to ancient human migrations. Mol Biol Evol. 2012;29(1):359–365

6. Hammer 2001: Hammer, Ø., Harper, D.A.T., Ryan, P.D. 2001-2018. PAST: Paleontological statistics software package for education and data analysis. Palaeontologia Electronica 4(1): 9pp., http://palaeo-electronica.org/2001_1/past/issue1_01.htm, Natural History Museum, University of Oslo

7. Istvanovits 2017: Eszter Istvanovits, Valeria Kulcsar. Sarmatians – History and Archaeology of a Forgotten people. Maintz 2017.

8. Karachanak et al 2013: Sena Karachanak, Viola Grugni, Simona Fornarino, Desislava Nesheva, Nadia Al-Zahery, Vincenza Battaglia, Valeria Carossa, Yordan Yordanov, Antonio Torroni, Angel S. Galabov, Draga Toncheva, and Ornella Semino. Y-Chromosome Diversity in Modern Bulgarians: New Clues about Their Ancestry. https://www.ncbi.nlm.nih.gov/pmc/articles/PMC3590186/?fbclid=IwAR3jzec8zI4iiagoE7zmPe63ZYHnwhCTHMPsG7_VokDCGA8ObZz1qhepDms#pone.0056779-Dobrev1. Taken on 10.05.2019

9. Petar Golijski 2006: Balgarite v Kavkaz I Armenia (II-X c.), Sofia 2006.

10. Beshevliev 1981: Веселин Бешевлиев. Първобългарите. Бит и култура. София, 1981. Veselin Beshevliev. Parvobalgarite. Bit i kultura. Sofia, 1981.

11. Botalov 2010: С.Г. Боталов. раннетюркские ареапы восточной европы в системе кочевой цивипизации евразии. В: Купьтуры евразийских степей второй поповины I тысячепетия н.э.вопросы межэтнических контактов и межкупьтурного взаимодействия, Самара 2010, 4–41. S.G. Botalov. Rannetjurkskie arealy vostochnoj evropy v sisteme kochevoj civilizacii evrazii. In: Kul’tury evrazijskih stepej vtoroj poloviny I tysjacheletija n.je.voprosy mezhjetnicheskih kontaktov i mezhkul’turnogo vzaimodejstvija, Samara 2010, 4-41.

12. Vaklinov 1977: Станчо Вакпинов. Формиране на старобъпгарската куптура. София, 1977. Stancho Vaklinov. Formirane na starobalgarskata kultura. Sofia, 1977.

13. Gavrituhin 2003: И.О. Гавритухин, А.В. Пьянков. Крым, Северо-Восточное Причерноморье и Закавказье в эпоху средневековья. IV-XIII века, 186–199. Москва, 2003. I.O. Gavrituhin, A.V.P’jankov. Krym, Severo-Vostochnoe Prichernomor’e i Zakavkaz’e v jepohu srednevekov’ja. IV-XIII veka, 186-199. Moskva, 2003.

14. Komatarova 2011: Е. Коматарова-Балинова. Обредът на трупоизгарянето в некрополите на Салтово-Маяцката култура и биритуалните некрополи на Долен Дунав. В: Салтово-маяцькаарх археологічна культура: 110 років від початку вивчення на Харківщині. Харков, 2011. E.Komatarova-Balinova. Obredat na trupoizgaryaneto v nekropolite na Saltovo-Mayatskata kultura i biritualnite nekropoli na Dolen Dunav. V: Saltovo-mayatsykaarh arheologichna kulytura: 110 rokiv vid pochatku vivchennya na Harkivshtini. Harkov, 2011.

15. Komatarova 2012: Е. Коматарова-Балинова. Сравнительная характеристика элементов погребального обряда биритуальных могильников нижнедунайского населения и салтово-маяцкой культуры. В: Степи Европы в эпоху средневековья. Т.9. Донецк, 2012, 135–165. E.Komatarova-Balinova. Sravnitel’naja harakteristika jelementov pogrebal’nogo obrjada biritual’nyh mogil’nikov nizhnedunajskogo naselenija i saltovo-majackoj kul’tury. V: Stepi Evropy v jepohu srednevekov’ja. T.9. Doneck, 2012, 135-165.

16. Komatarova 2013: Е. Коматарова-Балинова. Хокери и псевдохокери от биритуалните некрополи в североизточна България: Възможности за интерпретации. В: Салтово-маяцька археологічна культура: проблеми та дослідження. Харков 2013, 82–88. E.Komatarova-Balinova. Hokeri i psevdohokeri ot biritualnite nekropoli v severoiztochna Bulgaria: Vazmozhnosti za interpretatsii. V: Saltovo-mayatsyka arheologichna kulytura: problemi ta doslidzhennya. Harkov 2013, 82-88.

17. Kyzlasov 1991: Игор Кызласов. Новые данные о произхождении и распространении древнетюркской рунической письмености Евразии. В: Проблеми на прабългарската история и култура. София, 1989,16–27. Igor Kyzlasov. Novye dannye o proizhozhdenii i rasprostranenii drevnetjurkskoj runicheskoj pis’menosti Evrazii. In: Problemi na prabalgarskata istorija i kultura. Sofia, 1989,16-27.

18. Marchenko 1996: И.И. Марченко. Сираки Кубани (По материалам курганых погребений Нижней Кубани.) Краснодар 1996. I.I.Marchenko. Siraki Kubani (Po materialam kurganyh pogrebenij Nizhnej Kubani.) Krasnodar 1996.

19. Minaeva 1951: Минаева Т.М. Археологические памятники на реке Гиляч в верховьях Кубани. В: «Материалы и исследования по археологии СССР» № 23. М.—Л., 1951. Minaeva T.M. Arheologicheskie pamjatniki na reke Giljach v verhov’jah Kubani. «Materialy i issledovanija po arheologii SSSR» № 23. M.—L., 1951.

20. Minaeva 1982: Т.М. Минаева. Раскопки святилища и могильника возле городища Гиляч в 1965 г. Москва, 1982. T.M. Minaeva. Raskopki svjatilishha i mogil’nika vozle gorodishha Giljach v 1965 g. Moskva, 1982.

21. Rashev 1991: Рашо Рашев. Две групи прабългари и две прабългарски култури. В: Историко-археологически изследвания. В памет на проф. Станчо Ваклинов, 29–33. Вепико Търново 1991. Rasho Rashev. Dve grupi prabalgari i dve prabalgarski kulturi. In: Istoriko-arheologicheski izsledvania. V pamet na prof. Stancho Vaklinov, 29-33. Veliko Tarnovo 1991.

22. Rashev 1992: Рашо Рашев. Запроизходанапрабългарите. В: Studia protobulgarica mediaevalia europensia. В чест на професор Веселин Бешевлиев, 23–33 Вепико Търново, 1992. Rasho Rashev. Dve grupi prabalgari i dve prabalgarski kulturi. In: Istoriko-arheologicheski izsledvania. V pamet na prof. Stancho Vaklinov, 29-33. Veliko Tarnovo 1991.

23. Rukavishnikova et al 2018: И. В. Рукавишникова, М. Ю. Меньшиков, И. И. Воробьев. Исследования нового биритуального могильника хазарского времени на северо-западном Кавказе. Кавказ в системе культурных связей Евразии в древности и средневековье. Кавказ в системе купьтурных связей Евразии в древности и средневековье, 361–364. I. V. Rukavishnikova, M. Ju. Men’shikov, I. I. Vorob’ev. Issledovanija novogo biritual’nogo mogil’nika hazarskogo vremeni na severo-zapadnom Kavkaze. Kavkaz v sisteme kul’turnyh svjazej Evrazii v drevnosti i srednevekov’e, 361-364.

24. Rusanova 1976: И. П. Русанова. Славянские древности VI-VII вв. Москва. 1976. I. P. Rusanova. Slavjanskie drevnosti VI-VII vv. Moskva 1976.

25. Stanchev 1958: Станчо Станчев (Вакпинов). Некропопът при Нови пазар. София, 1958. Stancho Stanchev (Vaklinov). Nekropolat pri Novi pazar. Sofia, 1958.

26. Suhanov 2018: Е. В. Суханов, А. Н. Свиридов. Новые раннесредневековые погребения с территории западного предкавказья. Росийская археопогия, 2018, № 4, с. 114–129. E. V. Suhanov, A. N. Sviridov. Novye rannesrednevekovye pogrebenija s territorii zapadnogo predkavkaz’ja. Rosijskaja arheologija, 2018, № 4, s. 114–129.

27. Adams 2014: Noel Adams. Bright light in the dark ages. London 2014.

28. Csaky et al 2019: Veronika Csáky, Dániel Gerber, István Koncz, Gergely Csiky, Balázs G. Mende, Bea Szeifert, Balázs Egyed, Horolma Pamjav, Antónia Marcsik, Erika Molnár, György Pálfi, András Gulyás, Bernadett Kovacsóczy, Gabriella M. Lezsák, Gábor Lőrinczy, Anna Szécsényi-Nagy, Tivadar Vida. Genetic insights into the social organization of the Avar period elite in the 7th century AD Carpathian Basin. https://www.biorxiv.org/content/10.1101/415760v2 taken at 10.05.2019

29. Karatay 2009: Osman Karatay. The Bulgars in Transoxiana: Some Inferences From Early Islamic Sources, Migracijske i Etničke Teme, XXV/1-2 (Yaz 2009), s.69–88.

30. Nesheva et al 2015: D. V. Nesheva, S. Karachanak-Yankova, M. Lari, Y. Yordanov, A. Galabov, D. Caramelli, D. Toncheva. Mitochondrial DNA Suggests a Western Eurasian Origin for Ancient (Proto-) Bulgarians. Human Biology, 87(1):19–28. 2015.

31. Rapp 2003: Rapp, Stephen H. Studies In Medieval Georgian Historiography: Early Texts And Eurasian Contexts. Peeters Bvba, 2003.

