## Supplement for "Genetic evidence suggests relationship between contemporary Bulgarian population and Iron Age steppe dwellers from Pontic-Caspian steppe"

### Archaeological overview on the formation of Asparukh’s Protobulgarians

Todor Chobanov Ph.D.,Ass.prof., Svetoslav Stamov MA, Duke University

Todor Chobanov is an archaeologist. His interests include Protobulgarian origins, Pagan culture of Danube Bulgaria, Medieval architecture
Svetoslav Stamov MA is a cultural and physical anthropologist, who graduated in Duke University, USA. His various interests include population genetics, Protobulgarians and others.

Summary

The present study aims to properly introduce the newly acquired genetic data from various surveys into the debate for the origin of the Protobulgarians. A part of the research focuses on the archaeological definition of Protobulgarians with all its key features, finally identifying the most likely key area of ethno genesis, starting from 2nd c. BC – the Kuban river area. The archaeological features that prove solid contacts with early Alans are discussed, as the emerging of biritualism or even multiritualism in the same zone. Finally, the available genetic data is processed with the Past software to produce principle component analysis (PCA) for the modern Bulgarians, comparing them with various ancient populations. The results prove close ties with Saltovo-Mayaki peoples and particularly with Caucasian Alans. Based on this observations and particularly the placing of various Alan samples firmly within the neighbor joining tree of modern Bulgarians, the enthnogenesis zone is reviewed again and the early start – 1st-2nd c. is confirmed. The general conclusion is that Protobulgarians were a mixture of Late Sarmatians and older Caucasus populations, closely related to the Alans and preserving their genetic inheritance even after arriving on the Balkans and mixing with Slav peoples and the remnants of the local Late Antiquity peoples.

The present day country of Bulgaria, often designated as “Danube Bulgaria” in various archaeological and historical works, was established during the centuries following the Hun period in Europe and undoubtedly represents one of the most enduring results of the Migration period in Europe (Völkerwanderung in German). The identity and origin of the people who stood behind this enduring act of state-creation still remains unclear and often – hotly debated.

The historical, archaeological, onomastic and linguistic research that started in 18th c. allowed scientists both domestic and foreign to reach certain conclusion grouped in around several major theories. The most comprehensive summary on the matter up the late 20th c. period was delivered by Veselin Beshevliev. He would then summarize that there were in the past four major theories about the origin of the Protobulgarians – labeled by him as “Tracian”, “Slavic”, “Finish” and “Ural-Altaic” (**Beshevliev 1981**, 15). Beshevliev carefully tried to distinguish between “Hunnic origin” views and the Ural-Altaic theory. He would also conclude that “The Danube Bulgarians are descendants of the Bulgarian tribe Unogundurs - “that has no clear ethnic identification, but was undoubtedly of Turkic origin” (**Beshevliev 1981**, 20). However, he realized based mostly on available linguistic evidence and especially on tribal and personal aristocratic names, that the Protobulgarians before Danube Bulgaria were not a homogenous group and different identities were in the process of mixing at least on aristocratic level. Beshevliev’s views demonstrated the level that the debate about the origin of Protobulgarians has reached in the late 20th c., but the situation was about to change.

The main agent of this change of views in Bulgarian archaeology was the prominent Bulgarian scientist Rasho Rashev (1943-2008). Rashev supported Beshevliev’s view that the Protobulgarians were not a homogenous group and different suffixes attached to tribal and clan names actually represent different ethnicities – “-ar”, “-gir” and “-dur” for different Turkic, Ugrian and Iranian elements, that probably merged together during the Hun era of Europe (**Rashev 1991**, 29). He then concludes that the 12 years calendar used by the Protobulgarian elite was probably Turkic, similar to the ones used in Middle Asia. This was the result of events in the 6th century, when the different Protobulgarian tribes in the Caucasus area fell under the control of the Western Turk khaganate. The key finds related to those elites are the Pereschepina complex and the Voznesenka memorial complex, but those examples could not be related to the culture of the “New lands” (Danube Bulgaria) with the exception of grave 3 mound 5 from Madara (**Rashev 1991**,30). Rashev then examines the ordinary people. Even back in the early 90ties about 20 pagan period necropolises – with cremations, with inhumations and mixed - were known. He concluded that the inhumations, which represented about one third of the total number of graves, show direct parallels with "Sarmatian burial practices from the 1st-2nd century" in the region between Danube and Don rivers (**Rashev 1991**, 31). The features in question are the Northern or Western orientation of skeletons, the ash layers on the bottom or in the fill of the grave pit, the post-mortem destruction of the skeleton, brachycranic types and artificial cranial deformation. As for the cremations, Rashev proposes that the ones in biritual necropolises could have been Protobulgarian but not Slavic (**Rashev 1991**, 31). He also underlines that very few Turkic loans exist into the well-studied Bulgarian language from the 10th century, which means that the Turkic speakers amongst the elite were very few in number. As for the Iranian language of the regular population, he proposes that in areas as the forest-steppe of middle Dnieper river the Sarmatian Protobulgarians entered into prolonged contacts with the Slavs, leading to their early ethnolinguistic Slavicization. The overall solution of the problem for the ethnicity of the Protobulgarians offered by Rashev is the same as the one proposed by Stancho Vaklinov after the excavation of the Novi Pazar necropolis and by Simirnov, Artamonov and others in relation to Volga Bulgarians and the genesis of the Saltovo culture – that a tiny Turkic elite ruled over host of Sarmatians and Ugric peoples (**Rashev 1992**, 26).

The link in the chain between 3rd-4th c. Late Sarmatians and 7th c. Protobulgarians was pointed by Rashev to be the 6th -7th c. Penkovsk culture, that inherited the traditions of the preceding 5th century Kievska culture from the middle Dnieper area. The later have likely included Chernyakhov peoples and traditions, as later did Penkovsk culture. It conveniently spread in the forest-steppe zone around the left tributaries of Dnieper west to Seret where it contacted the Slavic Praga-Karchak culture. The Penkovsk culture practiced cremations and disappeared in late 7th-early 8th century, in the eve of the Khazar era. Interestingly, in the south Penkovsk culture reached the areal of the "Sivashovka group", presently labeled as "Syvashovka horizon" and thus – the area of inhumation practitioners. Contacts apparently influenced the material culture - the pottery of Penkovsk culture included different types – alongside hand-made biconical pots there were also wheel-produced round pots from gray clay decorated with indented shining band areas. The later and very interesting type of pottery, was widely designated after "Pastirsk(oe)" fortress, where many examples where discovered, as Pastirsk type (**Rashev 1992**,27-29). Rashev built his theory on the works of Slavic history researcher Rusanova, who established that all Penkovsk pottery types, including the Pastirsk types, have direct Chernyakhov predecessors and that there was dominating presence of non-Turkic (Iranian) peoples in Penkovsk culture. The Antes proper were direct descendants of the old Sarmatian population of the region who in 6th-7th c. were mixing with arriving Slavic peoples. The typical biconical pots are not Slavic, are missing in the Praga-Karchak culture and there is a group of hand-made pots related to Sarmatian, “Avar” and Saltovo types. The Slavic arrivals were gradually passing their languages to the Antes, who were Slavicized linguistically (**Rusanova 1976**, 75-112). This theory enabled Rashev to conclude that in his westbound movement Asparukh, supposedly starting his trip from the inner territories of Great Bulgaria east of Azov Sea, stimulated the Penkovsk population to move with him to the emerging Danube Bulgaria, which is a very sound explanation about the multiple finds of Pastirsk type pottery there in the 7th-9th c. period. This actually represents a second and later group of finds with this type pottery on the Lower Danube the previous one being 6th-7th c. followed by a pause. It also explains the quick language slavicization of the new state in opposition to the missing traces of Turkic language speakers. “The non-Slavic population of Danube Bulgaria were the descendants of the old Iranian peoples of Eastern Europe (**Rashev 1992**,28-30).

In recent years, it is worth mentioning the attempt to redefine “Early Turkic areas” in Eastern Europe by Sergei Botalov. He claims that the “*Huno-Bulgarian*” area was established as early as mid 3rd century AD in the steppe zones next to Northwest Caucasus (Kuban river area). As earliest sites he points at Yúzhnaya Ozeréyevka and Tzemdolinsk necropolises, as well as other sites near Novorosiisk and Anapa. The defining features are the earliest known cases with western orientation of the heads of the deceased buried with horses or cases of separate horse burials. Those horse finds in human graves or separate horse graves are today considered the earliest of this type in Eastern Europe (**Botalov 2010,11**). The “*Huno-Bulgarian*” area stretched south to the coastal area of Tuapse, with the Bzid 1 necropolis. The inventories of the graves are typical for the Late Sarmatians, especially same period finds from the Volga and Don rivers. (**Botalov 2010**, 4; **Gavrituhin 2003**). Botalov concludes that this “*Huno-Bulgarian*” area concluded its formation in the Hunnic and Post-Hunnic period (5th-6th c.) and continued its existence up to 8th century. The inhumation ritual prevailed over cremation, which became dominant only after 8th century. The graves are simple rectangular pits with stone plates tiling or stone cassettes inside the grave, head orientation is mostly western, but eastern is also present. In many necropolises, most notable being Durso and Bzid 1, there are horse burials. Botalov considers that the core of the area was in its northern part (closer to the Steppes?) where present day Tamanski and Novorosiiski districts are and where the key necropolises Durso and Borisovski are situated. Notable Russian researchers Gavrituhin and Pjankov connect this group with Kuvrat’s Great Bulgaria (**Gavrituhin 2003**,192).

In the south, the Protobulgarian area stretched to Tuapse and Sochi, in the east - the upper Kuban river area, where the *Giliach* necropolis with rectangular pits with stone plating and east or north head orientation and typical "Hun and post-Hun" inventories was discovered. The *Giliach* necropolis includes pottery with strong Central and Eastern Caucasus influence, but east from it begins another area, which is dominated by mound or catacomb burials (**Botalov 2010**, 4-5; **Minaeva 1951**; **Minaeva 1982**). Defined as described above, the "*Huno-Bulgarian*" area served according Botalov as "cradle" for the much greater "*Northern Black sea area*" which shaped in two major stages – 6$^{th}$-7$^{th}$ century – the political period of Western Turkic khaganate domination and Great Bulgaria supremacy. This greater area later concluded its existence by gradual inclusion into Danube Bulgaria and the Khazar khaganate (late 7$^{th}$ -8$^{th}$ c.). This "*Northern Black sea area*" includes notable necropolises as Netailovsk, Krimski, Krasnaja Gorka in its eastern section; Novi Pazar, Kulevcha, Galiche, Dolni Lukovit I, Topola, Istritza and others – in its western section (**Botalov 2010**, 10-11).

According to Botalov the foundation of the "*Huno-Bulgarian*" area was established by "the Bulgarian population" of the North-eastern Black sea area – tribes as Utigurs, Kutrigurs, Saragurs, Onogurs and etc. that came from " Western Altai and Central Asia" during the Hun age and most likely – during Atila's wars. Those tribes mixed with the post-Hun Sarmatian-Alan population that delivered to it cultural features that were "long kept" by early medieval Bulgarians – pits with niches, northern orientation, artificial cranial deformation. Interestingly, Botalov is also willing to support Stancho Vaklinov's conclusions formulated after the excavation of Novi Pazar necropolis that "*the foundation of of Protobulgarian culture is rooted in Late Antiquity Sarmatian-Alan culture*" (**Stanchev 1958**, 112-116; **Botalov 2010**,11). The final accord of Protobulgarian formation, concluded Botalov, came with the arrival of Avars and Sabirs and, finally, of the Ashina Turks, who concluded subjugating treaties with the two major groups – Kutrigurs and Utigurs (in 575 and 582).

The abovementioned area demonstrates one very important archaeological feature. Apparently birirualism on the Lower Danube has its closest parallel in various *Saltovo* necropolises like Krasnaia gorka (Harkov area). One very specific common feature is the use of raw or roughly shaped stones to build the grave (**Komatarova 2012**,159-160). Statistical analysis of the five variants of grave directions demonstrate that the dominant one for the Lower Danube necropolises was the northern one with deviations to east and west. The situation in *Saltovo* necropolises proper is somewhat different - the Northern/North-East orientation is dominant only in biritual placed in Criema and Taman peninsula area (**Komatarova 2012**,153). The South-Southwest orientation that is predominant for the

*Zlivkinski* type cemeteries is completely missing on the Lower Danube. The general conclusion could be that the Protobulgarians who separated from the proto-Saltovo massive (Great Bulgaria period) and who arrived with Asparukh on the Lower Danube settled early (late $7^{th}$ c.) were moving out from the abovementioned areas of the Crimea and Taman peninsula and not from the Harkov area to the North. The "classic" Saltovo period started 70 years later – around mid $8^{th}$ c. (**Komatarova 2012**,160). The early separation (after 650-660) and quick settling down (after 680) could be the explanation for the stronger preservation of some Late Sarmatian features on the Lower Danube, but it also could derive from a specific regional traditions. This northern orientation of graves in northwest Caucasus area could be explained by traditions deriving from "post-Maeotian population". This observation is confirmed by the newest discoveries of sites in the same area – as the $5^{th}$-$7^{th}$ c. Varnavinskoe-3 site (Krasnodar area) where the northern orientation of the inhumations is compared by the researchers with the Borisovski and Durso necropolises (**Suhanov 2018**,125).

One very interesting feature of cremation burials on the Lower Danube is the use of stone or brick cassettes. This grave setting is most common in the necropolises close to the Black sea – Devnja 1 and Devnja 3, Varna, Balchik, Topola, Bdintzi and Kapul Viilor. In the *Saltovo* culture areas, stone cassettes are most common in the Kuban-Black sea area – especially in the graveyard Borisovski, Krasnodar area (77.5% of the cremation graves), where the cassettes have no bottoms and are prepared with stone slabs only (**Komatarova 2011**,34). This technology for the preparation of the stone cassettes finds its closest parallel in the cassettes of early Alans in the Central Caucasus and Kuban river area.

The use of urns for cremations in biritual necropolises attested on the Lower Danube is also a common feature in Saltovo sites – more dominant in Donetzk area (Suhaja Gomolsha site) and far less common in the Kuban area sites – Borisovski, Durso and Kazanovo-2. While on the Lower Danube cremations are attributed both to the Slavic populations and Bulgarians proper in the northwest Caucasus area they must be attributed to non-Slavic groups, possibly indigenous people of the Caucasus or other cremation practitioners, living alongside the Protobulgarians (**Komatarova 2011**,35-36). The emerging biritualism during $4^{th}$-$7^{th}$ c. in the northwest Caucasus area is the most plausible source for the biritualism on the Lower Danube. It is notable that biritualism in the northwest Caucasus area continued to exist in the "classic" *Saltovo* area, as attested by the newest find of a $9^{th}$-$10^{th}$ c. site in the Krasnodar area in which cremations are dominant – 15 out of 19 graves and include both cremations in urns and in pits. The researchers relate it to the fourth period of the Durso site (**Rukavishnikova et al 2018**,364).

One more feature intrigued the researchers in Bulgaria and elsewhere emerging from excavations of biritual necropolises on the lower Danube – Novi Pazar, Topola, Devnja-1, Devnja-3, Bdintzi, Balchik. About 30 burials are cases of hoker or “pseudo-hoker” with typically shallow grave pits and lack of grave goods are known on the Lower Danube, sometimes reaching 12-17 % of the total number of graves (Novi Pazar being the “champion” with 17%). There is not any evident correlation between the inhumation-cremation ratios (**Komatarova 2013**, 83). It is very important to notice that the phenomenon is not at all registered in the many Bulgar-associated graves before the Saltovo period. The absence of such burials in the Pokrovsk and Sivashovka periods and its emerging in both Saltovo and Danube Bulgaria necropolises demands a plausible explanation. It was delivered by Evgenia Komatarova based on previous analysis – “alien ethnic elements” of Alan-Sarmat nature influenced the Bulgars. Komatarova highlighted the case of Dimitrievski necropolis where amongst the dominant (Alan) catacomb burials there are nine cases of hokers, others are registered in necropolises like Majatzki and Lubljanski. More intriguingly, Lower Danube hokers from Devnja and Varna that have two or three individuals show notable similarities with the catacomb burial ritual(Komatarova 2013,86). Another key fact registered for the Lower Danube hokers is that some of them (Topola) contain individuals with artificial cranial deformation (**Komatarova 2013**, 86-87).

The developments in the field of Archaeology, related to Protobulgarians, could be successfully supplemented with data from other sources. In the last years, genetic studies have firmly entered the arsenal of Archaeology with lasting consequences. One of the first projects, involving data from present day Bulgaria, was published in 2013. It was a major representative survey of present dale male lineages in Bulgaria (over 800 individuals) and it revealed that “Haplogroups C, N and Q, distinctive for Altaic and Central Asian Turkic-speaking populations, occur at the negligible frequency of only 1.5%.” (**Karachanak et al 2013**). This major observation was somewhat a surprise considering the supposed Turkic-Altaic origins of Protobulgarians, the serious impact of 11th-12th century late nomads as Pechenegs , Cumans and Uzi peoples and the five centuries of Ottoman rule. The authors of the survey concluded that *“...our data suggest that a common paternal ancestry between the proto-Bulgarians and the Altaic and Central Asian Turkic-speaking populations either did not exist or was negligible...”*( **Karachanak et al 2013**, abstract). The later conclusion could mean that the Protobulgarians were really too few, or as shown by the present one and other DNA surveys – originally did not have Central (Inner) Asian ancestry at all.

The next major project that is relevant to the topic of this article was published in 2015. It was the first ever mtDNA sampling of materials from 3 8th-9th c. necropolises on the Lower Danube (Ancient DNA). Again, The main haplogroup H (H, H1, H5, and H13) prevalent in European populations has a 41.9% frequency in modern Bulgarians, and it was observed in 7 of 13 proto-Bulgarian samples. The researching team found no evidence of East Asian (F, B, P, A, S, O, Y, or M derivative) haplogroups. Their overall conclusion was that *"…Our results suggest a western Eurasian matrilineal origin for proto-Bulgarians, as well as a genetic similarity between proto- and modern Bulgarians*." (**Nesheva et al 2015**, 1).

However, a major picture demonstrating the complex realities in the Eurasian steps is necessary in order to properly analyze the results from the two abovementioned projects and attached them to more specific source area – a possible homeland for the formation of the Danube Protobulgarians. Such a picture came from a capital article that was published recently in the "Nature" magazine – "137 ancient human genomes from across the Eurasian steppes"(**137 genomes 2018**). This is the first massive whole genome capture project that contains various samples from the Eurasian steppes, that are compared amongst themselves and against existing populations. The genetic data from this survey was accompanied with a vast additional text (supplements), thus providing information about the origin and cultural context of each sample thus enabling us to establish patterns and more importantly – trace migration routes and mixing scenarios.

The global conclusions from the "137 genomes" survey successfully highlight the population dynamics in the Eurasian steppes. Significant ancestry from Neolithic farmers is well evident in Late Bronze Age steppe herders and Iron Age steppe nomads that is not shared with Yamna herders (137 genomes,370). The strongest flow of European Neolithic farmers is evident in samples from the Hungarian plains – "Hungarian Scythian". The percentage of East Asian genetic influence in Hun samples is relatively low – 10-15%, similar to the levels of East Asian influence in the preceding Scythian populations. The Western Huns around 2000 years ago show some Caucasian ancestry as well (even before entering Eastern Europe), but the percentage is even higher that amongst the preceding Scythians – about 30%. The general outline of haplogroups for the Western Huns is European with one exception, where the O haplogroup is attested (Chinese contacts?) but still the clear division between Western and Eastern Huns has been confirmed by the DNA data. The relation between the Western Huns and the preceding Saka(Scthytian)-Sarmathian groups in the steppes has been fully confirmed and no room remains for any alternative development. The survey also brings interesting results about the Early medieval Alans (6-9th century AD) who are generally accepted to be descendants of the late Sarmatian culture (ending around around 4th c. AD). The strong connection is confirmed, but it also becomes clear that when settling in the Caucasus area they started mixing with the local population. Evidently, at 4th-5th c. the local Caucasian population already has massively transformed them to the point that "their genetic ancestry was already indistinguishable from the neighboring Caucasian populations by the 5th-6th Century CE" (**137 genomes-supp,** 194-195)

By analyzing ancient DNA samples from Bronze Age, Iron Age and medieval Western and Central Eurasia, we try to establish the source population(s) and the timing of the additional Caucasian admixture in contemporary Bulgarians. We also review already published genetic research on the topic in the scientific literature and attempt to identify what is already known about the timing and the hypothesized source population. We also test several well-known historical hypothesizes about the origins of contemporary Bulgarians and early medieval Protobulgarians.

We use PAST Software for ancient DNA analysis and to plot over a hundred ancient DNA samples against the DNA samples of contemporary Bulgarians. We conduct principal component analysis, measure genetic distance between the selected samples and build a genetic based on the genetic clustering of the samples with several contemporary individuals from SE Europe from selected contemporary ethnicities, with focus on contemporary Bulgarians. All genetic trees and plots have been made with PAST software for palaeogenetic DNA analysis.

The testing produced important results enabling us to clarify the relationship between contemporary Bulgarians and ancient individuals from Western, Eastern and Central Eurasia. Our results do not support central Asian or East Asian origin of contemporary Bulgarians (Fig. 2, Fig. 3, Fig. 4, Fig. 5). Our results also do not support central Asian or far eastern origins of the early medieval Protobulgarians. (Fig. 2, Fig. 5) Instead, the results reveal massive Caucasian component in both contemporary Bulgarians and Medieval Protobulgarians, implying major contribution of the Protobulgarians to the contemporary Bulgarian genetic make-up and also implying at least partly Caucasian origins of the Protobulgarians. Using statistical genome-wide analysis, we detected nontrivial genetic connection between medieval Protobulgarians and the inhabitants of Bronze Age Armenian plateau as well as to Iron Age Sarmatians from Northern Caucasus. Our analysis also suggests surprising connection between contemporary Bulgarians and Iron Age Scythians from Hungarian plain.

We noted that SM (Protobulgarian-Alan) influence among contemporary Balkan nations has its strongest representation in contemporary Bulgarians (Fig 4) where it arrives directly and this Protobulgarian influence in the other Balkan nations is mediated by the contemporary Bulgarians who channel it.

Conclusions from the DNA data analysis are as follows: the results suggest that SM related populations are the precursor of contemporary Bulgarians. This makes SM culture at its precursor stage (600-700) a leading candidate for the source population of Asparukh Bulgarians. These results also suggest that Asparukh's tribe(s) are indistinguishable from the Sarmato-Alanic groups from Early middle ages and Late antiquity and, surprisingly, do not carry Siberian and Central Asian admixture on the Balkans with them, as even the Protothracians of early Bronze Age Balkans show bigger Siberian admixture (probably reflective of their IE ANE component). Unlike Protothracians and the early Slavs, Protobulgarians carry substantial Caucasus Mountain admixture, related to the tribes from Bronze Age Armenian plateau and seems to have transmitted this admixture to the contemporary Bulgarians (see Fig. 2). The relationship between Protobulgarians and Sarmato-Alanic tribes from the Late antiquity and Early medieval epoch remains to be clarified further, however genome wide-data suggest that Protobulgarians were themselves an admixture in equal proportions between two close, but distinct populations –1. Alano-Sarmatian tribe from the region north of Caucasus with some Kangju link to it and 2. An unknown tribe(s) from native Caucasian origin originating in what is now Armenian Plateau. Both Scythian samples from the Hungarian steppe and the Alans from Saltovo-Mayaki culture bear strong genetic resemblance to the Bronze Age Caucasian samples, which is missing in central Asian nomads but is presented in the contemporary Bulgarians (see fig 1, fig 2, fig 3 and fig 4).

Our results cast a serious doubt on a connection between Inner Asian nomadic tribes from Antiquity and the Protobulgarians-Alans from SM culture and Northern Caucasus. It is notable that recent sampling of Avar graves has proven beyond any doubt that significant part of the Avars was of Inner Asian origin and they kept marrying in certain closed circles for a long period ( **Csáky et al 2018**, passim). The surveys of the Avars continued by an extended team bringing even more conclusive results - the Principal Component Analysis (PCA) of the Avar dataset was compared with haplogroup frequencies of another 47 ancient groups, proving that the Avar elite shows affinities to Asian populations: close to 15th-19th centuries Yakuts from East Siberia and to two ancient populations from China. The Avars are firmly related to South Siberian Bronze Age populations and fall with Iron Age and medieval

Central and East-Central Asian sample sets of the Ward-type clustering tree (**Csaky et al 2019**, 6).

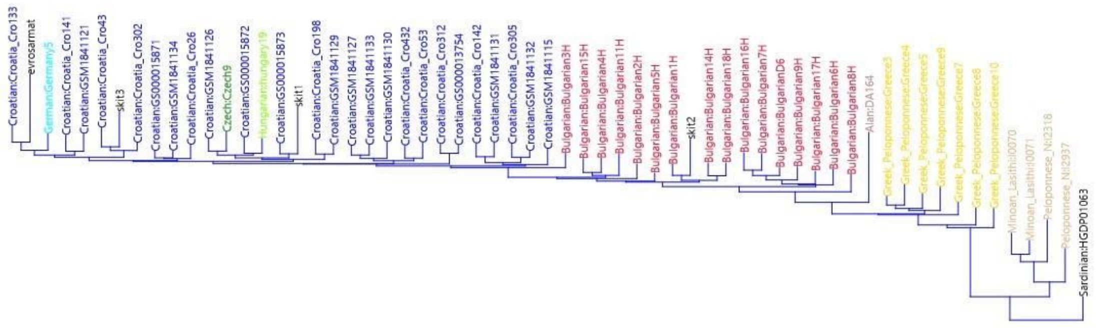


FIG 6 Ancient samples placing amongst modern peoples-1. Scythians from Hungarian plain and Alan samples fall firmly within the neighbor joining tree of present day Bulgarians, Scythians fall within the tree of modern Croatians suggesting strong Scythian component in the ethno genesis of Western Slavs. Plot: Sv.Stamov

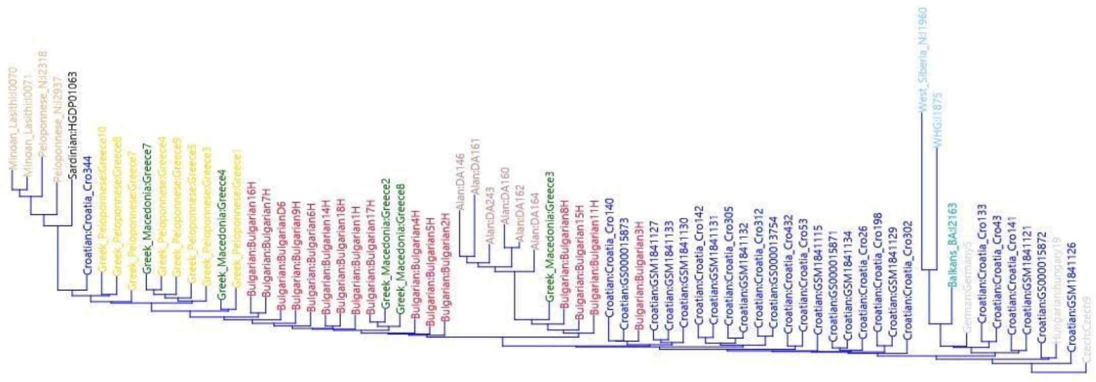

FIG 7 Ancient samples placing amongst modern peoples-2. The Various Alan samples from the Northern Caucasus fall firmly within the neighbor joining tree of contemporary Bulgarians. Plot: Sv.Stamov

We also clarified the origin of this Caucasian component further and managed to split the Caucasian component coming from SM from the Caucasian components already presented on the Balkans prior to Protobulgarian migration. We established that while all three carry somewhat similar Caucasian component, the signal, coming from SM is the strongest in contemporary Bulgarians, the signal coming from Bronze Age Thracians is the strongest in contemporary Croatians and the signal, coming from Bronze Age Minoans is the strongest in contemporary Greeks. These three signals clearly differ from each other and their source populations are clearly distinguishable. Yet all tree carry an excessive Caucasian component, suggesting non-local origins for all three of them and suggestive of at least three different migrations from the Caucasus Mountains to the Balkans. However, contemporary Bulgarians have received their Minoan component mostly through population exchange with Byzantium and their Bronze age Thracian component trough admixture/population exchange with early medieval Slavs and Croats. The signal that distinguished contemporary Bulgarians from the other Balkan nations is the unique signature of SM-Alan peoples, who appear amongst the direct precursors of contemporary Bulgarians.

One major conclusion from the genetic data is that the Bulgars were more numerous than the Slav tribes they intermixed with upon foundation of medieval Bulgaria. Another key observation is the absence of Inner Asia ancestry. Since both Early medieval and contemporary DNA samples failed to yield any connection between both early medieval Bulgars and contemporary Bulgarians, from one side, and both ancient and contemporary central Asian groups, from another side, but revealed strong genetic connection to both Sarmato-Alanic DNA samples from the Late antiquity and Early medieval era, as well as Sarmato-Alanic mediated connection to the contemporary Caucasian groups, our research suggest Caucasus mountains and especially its adjacent regions to the North as a homeland of Protobulgarians and as a place where their ethno genesis developed, which effectively renders Protobulgarians as an ancient group within the range of Sarmato-Alanic tribes from the late antiquity, eventually displaced on the Balkans and Eastern Europe by the migration of the Huns and later - the expansion of the Khazar khaganate.

It also becomes evident that the Slavic component of Bulgarian medieval and present day population developed on a solid West Scythian ancestry, which is evident also amongst other east Europeans particularly amongst modern Croatians (Fig. 4, Fig. 6, Fig. 7). The placing of samples enables us to theorize that the West Slavic presence on the Balkans was stronger that East Slavic one and the similarities between Praga-Karchak and Popina-Garvan pottery actually could be explained by belonging to the same archaeological culture/population.

It is evident that neither the Avar nor the Turkic political dominance over the Bulgar tribes left a significant genetic trace. The features of Avar DNA ancestry (inner Asia haplogroups) are missing completely in modern Bulgarians/Protobulgarians or groups associated with the Protobulgarians – like Alans or Saltovo peoples. The first ever solid study of Avar DNA enables us to establish the clear difference between Avars and Protobulgarians on one side but also to conclude that unlike Protobulgarians the Avars did not intermix with their Slavic subjsects (**Csáky et al 2018**, passim). The evidence that the soldiers of the Turkic Khaganate are genetically closer to East Asians than the preceding Huns (**137 genomes**,372) also removes the option that during the Turkic political dominance in 6th-7th c. the Turks proper intermixed with their Protobulgarian subjects in the Caucasus area. One sample from elite warrior however suggests that the Turks were not settling but were displacing (recruiting) people from East Europe. Those observations leave only one possible origin for the discussed Turkic elements in Protobulgarian culture – the words from the Stone inventories and the calendar terms – they were accepted under the political hegemony of the Khaganate in 6th-7th c., as Rashev proposed in the early 90ties. Regretfully no DNA data from rich Protobulgarian graves is available at present (for example the Kabiuk grave circa 700) and we could not check the existing theories that there were various ethnicities amongst the elite (Turks, Ugrians, Sarmatians).

The recent successes of scholars like Komatarova in tracing the roots of Lower Danube Protobulgarian biritualism in Krasnodar area and generally – in the northwest Ciscaucasia (where the Byzantine sources placed Old Great Bulgaria) combined with DNA data – enable us to narrow the zone of Protobulgarian ethno genesis similarly to what was done by Botalov using general archaeological observations only. The features of the genetic data - demonstrating strong Caucasian signal - require that the initial (preliminary) stage must be set early – as early as the arrival of the Siraci, Roxolani and Aorsi tribes (around 4th c. BC ). The Siraci are particularly interesting, because they settle the area under consideration and successfully mixed with the local Maeotians bringing the Sarmatisation of the later (**Marchenko 1996**,120).

The next major stage started with the arrival of the late Sarmatians in the area in 1st-2nd c. AD, bringing with them artefacts similar to those found in Volga and Don area. Apparently, the political turmoil this migration caused in the Easteuropean Steppes of the 1st-2nd c. AD forced some peceding Sarmatian tribes to move – some to the West, including the Danube delta area, others to the south, closer to the Caucasus Mountain, in less exposed Ciscaucasus areas. Part of this movement is detected through the disappearance of the early Sarmatians from the steppes extending on the right bank of the Kuban river and their reappearing in the river's middle reaches (**Istvanovits 2017**,151-152). The ethno genesis of Protobulgarians probably started in this very area in the late 2nd and early 3rd c. with this new wave of Sarmatians to enter the Ciscaucasia and starting to live alongside indigenous (older) tribal groups and be influenced by their culture. Similar reconstruction of the events in the mid 2nd c. AD is supported by O. Karatay, who sees the Bulgars as "merely one of several tribes that crossed the Volga in the so-called Sarmatic wave" and also as a part of the Sarmatian confederation that ruled for a vast territory in the Volga region (**Karatay 2009**, 82-83).

The next stage started with the arrival of the late Sarmatians in the area in 1st-2nd c. AD, bringing with them artefacts similar to those found in Volga and Don area. It is notable that this new wave of peoples was highlighted as related to Protobulgarians as early as mid 20th c. by key scholars like Nikolay Merpert (Merpert 1958, 586-588). It was likely this migration in the area that caused the political turbulence north of the Caucasus reported by Movses Khorenatsi, who claims that "…in the days of Arshak there was massive dismay in the chain of the Great Caucasian mountain in the lands of the Bulgharians and many of them separated and arrived in our land and lived long under the Kol…" (Horenatsi 1990, 67). It was proposed by recent research that those settlers who crossed the mountain and settled preserved their "Vanandur" ethnic labeling and probably some original features for a long period while living in the Upper Basean area that bore the name Vanand (Golijski 2006, 643-644). The migration happened most likely in the time of king Valarsh II (185-198) and was followed by less peaceful events – several hostile invasions by the different tribes living north of the mountain (Golijski 2006, 644-645).

The strong Caucasian DNA signal in Protobulgarians, Alans and present day Bulgarians supports the historical data about long interaction between the Protobulgarians and the indigenous people of the Caucasus, particularly Alans, Armenians and probably even Iberians. Alternative explanation would be that the signal was passed to Protobulgarians through the massive inclusion of Caucasian Alans (having strong Caucasus inheritance themselves) but such version is less likely considering the key differences between Protobulgarian and early Alan archaeological cultures.

All those observations and especially the archaeological ones bring the conclusion that the dominant tribe of the Great Bulgaria period – the Unogondurs, known as Nandor to ancient Hungarians and V-n-n-t-r in Josephus the Khagan's letter  and Perso-Arab sources had somewhat different ethno genesis within a larger group of similar tribes and were particularly under strong Alan-Caucasian influence, inhabiting the southernmost sections of a larger ethno-genesis area – closer to the mountain chain itself. Archaeological features of this Alan influence include black-grey polished pottery, stone cassettes, hokers and others.  It is notable that the Unogondurs left strong impression in Armenian sources – we should recognize them in Movses Khorenatsi's Vanandur and Anania Shirakatsi's Olhontor-Blkar. This line of analysis – that there were different Bulgar tribes that were developing somewhat separately from each other but in neighboring areas is also confirmed by the famous list of Pseudo-Zahariah Rhetor. It counts the Bulgars twice – once as settled peoples with cities and neighbouring the Alans – those being the Unogondurs in the Ciscaucasus area and second time as nomads, living in tents – those probably being the one living in the steppes of the Don area – notably the Kutrigurs.

Living in the southernmost reaches of the described ethno-genesis area in the northwest Ciscaucasus – around the middle Kuban river area and south of it (Map 1), the Unogondur-Bulgars emerged on the historical scene after the Turks weakened their grip on the region in the early 7th c. The decline and dispersing or change of identity of Utrigurs and Kutrigurs, highlighted by Agathias of Mirinea (Pohl 2018,32),  enabled the Unogondurs to unite both related Bulgar and other, not-related groups into the Old Great Bulgaria of Theophanes and Nicephorus and emerge as the Ούννογουνδούρων βουλγάρων from the 8th - 9th c. Byzantine sources .

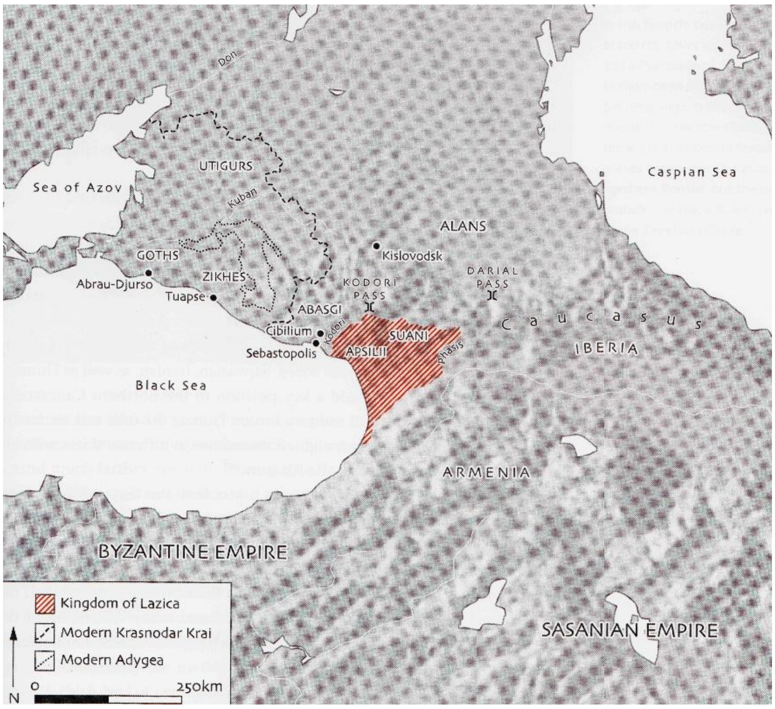


Map 1. The zone of Protobulgarian ethnogenesis in the Ciscaucasia. After **Adams 2014**,253

The research carried in this study, combining archaeological data and DNA research, brings the debate about the origin of Danube Protobulgarians into another level by identifying their Ciscaucasian "cradle" and thus – their likely Sarmatian-Caucasian origin, similar to those of early Caucasian Alans. It, however, could not answer many questions that will remain to be solved. The ethnolinguistic identity of the ancient Bulgars and particularly the channels of ancient Turkic influence need to be further clarified – was it only a political phenomenon of the 6th-7th c. or they were seriously mixing up earlier with indisputably early Turkic tribes like the Sabirs? How the ethnonym "Bulgar" appeared and when exactly, was it a political designation that spread over various tribes of different ethnocultural and ethnolinguistic identities? Was the Protobulgarian elite belonging to different ethnicities and what were they?

The future research – archaeological and genetic – will probably allow us to reexamine many of those issues, answer those questions and progress even further. But it is absolutely clear now that ancient DNA data will be invaluable asset for the proper understanding and interpretation of archaeological facts and – eventually – understanding History.

**References**

**Beshevliev 1981**: Веселин Бешевлиев. Първобългарите. Бит и култура. София, 1981. Veselin Beshevliev. Parvobalgarite. Bit i kultura. Sofia, 1981.

**Botalov 2010**: С.Г. Боталов. раннетюркские ареалы восточной европы в системе кочевой цивилизации евразии. В: Культуры евразийских степей второй половины I тысячелетия н.э.вопросы межэтнических контактов и межкультурного взаимодействия, Самара 2010, 4-41. S.G. Botalov. Rannetjurkskie arealy vostochnoj evropy v sisteme kochevoj civilizacii evrazii. In: Kul'tury evrazijskih stepej vtoroj poloviny I tysjacheletija n.je.voprosy mezhjetnicheskih kontaktov i mezhkul'turnogo vzaimodejstvija, Samara 2010, 4-41.

**Vaklinov 1977**: Станчо Ваклинов. Формиране на старобългарската култура. София, 1977. Stancho Vaklinov. Formirane na starobalgarskata kultura. Sofia, 1977.

**Gavrituhin 2003**: И.О. Гавритухин, А.В.Пьянков. Крым, Северо-Восточное Причерноморье и Закавказье в эпоху средневековья. IV-XIII века, 186-199. Москва, 2003. I.O. Gavrituhin, A.V.P'jankov. Krym, Severo-Vostochnoe Prichernomor'e i Zakavkaz'e v jepohu srednevekov'ja. IV-XIII veka, 186-199. Moskva, 2003.

**Golijski 2006**: Balgarite v Kavkaz i Armenia (II-X c.), Sofia 2006.

**Komatarova 2011**: Е.Коматарова-Балинова. Обредът на трупоизгарянето в некрополите на Салтово-Маяцката култура и биритуалните некрополи на Долен Дунав. В: Салтово-маяцькаарх археологічна культура: 110 років від початку вивчення на Харківщині. Харков, 2011. E.Komatarova-Balinova. Obredat na trupoizgaryaneto v nekropolite na Saltovo-Mayatskata kultura i biritualnite nekropoli na Dolen Dunav. V: Saltovo-mayatsykaarh arheologichna kulytura: 110 rokiv vid pochatku vivchennya na Harkivshtini. Harkov, 2011.

**Komatarova 2012**: Е.Коматарова-Балинова. Сравнительная характеристика элементов погребального обряда биритуальных могильников нижнедунайского населения и салтово-маяцкой культуры. В: Степи Европы в эпоху средневековья. Т.9. Донецк, 2012, 135-165. E.Komatarova-Balinova. Sravnitel'naja harakteristika jelementov pogrebal'nogo obrjada biritual'nyh mogil'nikov nizhnedunajskogo naselenija i saltovo-majackoj kul'tury. V: Stepi Evropy v jepohu srednevekov'ja. T.9. Doneck, 2012, 135-165.

**Komatarova 2013**: Е.Коматарова-Балинова. Хокери и псевдохокери от биритуалните некрополи в североизточна България: Възможности за интерпретации. В: Салтово-маяцька археологічна культура: проблеми та дослідження. Харков 2013, 82-88.

E.Komatarova-Balinova. Hokeri i psevdohokeri ot biritualnite nekropoli v severoiztochna Bulgaria: Vazmozhnosti za interpretatsii. V: Saltovo-mayatsyka arheologichna kulytura: problemi ta doslidzhennya. Harkov 2013, 82-88.

**Kyzlasov 1991**: Игор Кызласов. Новые данные о произхождении и распространении древнетюркской рунической письмености Евразии. В: Проблеми на прабългарската история и култура. София, 1989,16-27. Igor Kyzlasov. Novye dannye o proizhozhdenii i rasprostranenii drevnetjurkskoj runicheskoj pis'menosti Evrazii. In: Problemi na prabalgarskata istorija i kultura. Sofia, 1989,16-27.

**Marchenko 1996:** И.И.Марченко. Сираки Кубани (По материалам курганых погребений Нижней Кубани.) Краснодар 1996. I.I.Marchenko. Siraki Kubani (Po materialam kurganyh pogrebenij Nizhnej Kubani.) Krasnodar 1996.

**Minaeva 1951**: Минаева Т.М. Археологические памятники на реке Гиляч в верховьях Кубани. В: «Материалы и исследования по археологии СССР» № 23. М.—Л., 1951. Minaeva T.M. Arheologicheskie pamjatniki na reke Giljach v verhov'jah Kubani. «Materialy i issledovanija po arheologii SSSR» № 23. M.—L., 1951.

**Minaeva 1982**: Т.М. Минаева. Раскопки святилища и могильника возле городища Гиляч в 1965 г. Москва, 1982. T.M. Minaeva. Raskopki svjatilishha i mogil'nika vozle gorodishha Giljach v 1965 g. Moskva, 1982.

**Rashev 1991**: Рашо Рашев. Две групи прабългари и две прабългарски култури. В: Историко-археологически изследвания. В памет на проф. Станчо Ваклинов, 29-33. Велико Търново 1991. Rasho Rashev. Dve grupi prabalgari i dve prabalgarski kulturi. In: Istoriko-arheologicheski izsledvania. V pamet na prof. Stancho Vaklinov, 29-33. Veliko Tarnovo 1991.

**Rashev 1992**: Рашо Рашев. За произхода на прабългарите. В: Studia protobulgarica mediaevalia europensia. В чест на професор Веселин Бешевлиев, 23-33 Велико Търново, 1992. Rashev 1992: Rasho Rashev. Za proizhoda na prabalgarite. V: Studia protobulgarica mediaevalia europensia. V chest na profesor Veselin Beshevliev, 23-33 Veliko Tarnovo, 1992.

**Rukavishnikova et al 2018**: И. В. Рукавишникова, М. Ю. Меньшиков, И. И. Воробьев. Исследования нового биритуального могильника хазарского времени на северо-западном Кавказе. Кавказ в системе культурных связей Евразии в древности и средневековье, 361-364. I. V. Rukavishnikova, M. Ju. Men'shikov, I. I. Vorob'ev.

Issledovanija novogo biritual'nogo mogil'nika hazarskogo vremeni na severo-zapadnom Kavkaze. Kavkaz v sisteme kul'turnyh svjazej Evrazii v drevnosti i srednevekov'e, 361-364.

**Rusanova 1976**: И. П. Русанова. Славянские древности VI-VII вв. Москва. 1976. I. P. Rusanova. Slavjanskie drevnosti VI-VII vv. Moskva 1976.

**Stanchev 1958**: Станчо Станчев (Ваклинов). Некрополът при Нови пазар. София, 1958. Stancho Stanchev (Vaklinov). Nekropolat pri Novi pazar. Sofia, 1958.

**Suhanov 2018**: Е. В. Суханов, А. Н. Свиридов. Новые раннесредневековые погребения с территории западного предкавказья. Росийская археология, 2018, № 4, с. 114–129. E. V. Suhanov, A. N. Sviridov. Novye rannesrednevekovye pogrebenija s territorii zapadnogo predkavkaz'ja. Rosijskaja arheologija, 2018, № 4, s. 114–129.

**137 genomes**: P. Damgaard et al , 137 ancient human genomes from across the Eurasian steppes, Nature Springer, May 9 2018 (Nature volume 557), pp369–374.

**137 genomes sup**: 137 genomes. Supplementary information. https://www.nature.com/articles/s41586-018-0094-2

**Adams 2014:** Noel Adams. Bright light in the dark ages. London 2014.

**Csaky et al 2019:** Veronika Csáky, Dániel Gerber, István Koncz, Gergely Csiky, Balázs G. Mende, Bea Szeifert, Balázs Egyed, Horolma Pamjav, Antónia Marcsik, Erika Molnár, György Pálfi, András Gulyás, Bernadett Kovacsóczy, Gabriella M. Lezsák, Gábor Lőrinczy, Anna Szécsényi-Nagy, Tivadar Vida. Genetic insights into the social organization of the Avar period elite in the 7th century AD Carpathian Basin. https://www.biorxiv.org/content/10.1101/415760v2 taken at 10.05.2019

**Haak et al 2015:** Haak W, Lazaridis I, Patterson N, et al. Massive migration from the steppe was a source for Indo-European languages in Europe. Nature. 2015;522(7555):207-11

**Hammer 2001:** Hammer, Ø., Harper, D.A.T., Ryan, P.D. 2001-2018. PAST: Paleontological statistics software package for education and data analysis. Palaeontologia Electronica 4(1): 9pp., http://palaeo-electronica.org/2001_1/past/issue1_01.htm, Natural History Museum, University of Oslo

**Istvanovits 2017**: Eszter Istvanovits, Valeria Kulcsar. Sarmatians – History and Archaeology of a Forgotten people. Maintz 2017.

**Karachanak et al 2013:** Sena Karachanak,Viola Grugni, Simona Fornarino, Desislava Nesheva, Nadia Al-Zahery, Vincenza Battaglia, Valeria Carossa, Yordan Yordanov, Antonio Torroni, Angel S. Galabov, Draga Toncheva, and Ornella Semino. Y-Chromosome Diversity in Modern Bulgarians: New Clues about Their Ancestry.

https://www.ncbi.nlm.nih.gov/pmc/articles/PMC3590186/?fbclid=IwAR3jzec8zI4iiagoE7zmPe63ZYHnwhCTHMPsG7_VokDCGA8ObZz1qhepDms#pone.0056779-Dobrev1. Taken on 10.05.2019

**Karatay 2009:** Osman Karatay. The Bulgars in Transoxiana: Some Inferences From Early Islamic Sources, Migracijske i Etničke Teme, XXV/1-2 (Yaz 2009), s.69-88.

**Lazardis et al 2014:** Lazaridis I, Patterson N, Mittnik A, et al. Ancient human genomes suggest three ancestral populations for present-day Europeans. Nature. 2014;513(7518):409-13

**Mathieson et al 2018:** Mathieson I, Alpaslan-Roodenberg S, Posth C, et al. The genomic history of southeastern Europe. Nature. 2018;555(7695):197-203.

**Nesheva et al 2015:** D. V. Nesheva, S. Karachanak-Yankova, M. Lari, Y. Yordanov, A. Galabov, D. Caramelli, D. Toncheva. Mitochondrial DNA Suggests a Western Eurasian Origin for Ancient (Proto-) Bulgarians. Human Biology, 87(1):19-28. 2015.

**Rapp 2003**: Rapp, Stephen H. Studies In Medieval Georgian Historiography: Early Texts And Eurasian Contexts. Peeters Bvba, 2003.

**Yunusbayev et al 2012:** Yunusbayev B, Metspalu M, Järve M, Kutuev I, Rootsi S, Metspalu E, et al. The Caucasus as an asymmetric semipermeable barrier to ancient human migrations. Mol Biol Evol. 2012;29(1):359–365
